# Structures of Native Doublet Microtubules from *Trichomonas vaginalis* Reveal Parasite-Specific Proteins as Potential Drug Targets

**DOI:** 10.1101/2024.06.11.598142

**Authors:** Alexander Stevens, Saarang Kashyap, Ethan H. Crofut, Shuqi E. Wang, Katherine A. Muratore, Patricia J. Johnson, Z. Hong Zhou

## Abstract

Doublet microtubules (DMTs) are flagellar components required for the protist *Trichomonas vaginalis* (*Tv*) to swim through the human genitourinary tract to cause trichomoniasis, the most common non-viral sexually transmitted disease. Lack of DMT structures has prevented structure-guided drug design to manage *Tv* infection. Here, we determined the cryo-EM structure of native *Tv-*DMTs, identifying 29 unique proteins, including 18 microtubule inner proteins and 9 microtubule outer proteins. While the A-tubule is simplistic compared to DMTs of other organisms, the B-tubule features specialized, parasite-specific proteins, like *Tv*FAP40 and *Tv*FAP35 that form filaments near the inner and outer junctions, respectively, to stabilize DMTs and enable *Tv* locomotion. Notably, a small molecule, assigned as IP6, is coordinated within a pocket of *Tv*FAP40 and has characteristics of a drug molecule. This first atomic model of the *Tv*-DMT highlights the diversity of eukaryotic motility machinery and provides a structural framework to inform the rational design of therapeutics.

## Introduction

*Trichomonas vaginalis* (*Tv*) is a flagellated, extracellular parasite of the human genitourinary tract and causative agent of trichomoniasis, the most common non-viral sexually transmitted infection (STI), with 250 million infections per annum and global prevalence over 3%^1–3^. *Tv* infection is linked to increased rates of preterm delivery and mortality, genitourinary cancers, and HIV transmission, with disproportionate impact on women in developing countries^1–5^.

Though the antibiotic metronidazole can be curative, its carcinogenicity concern, increasing metronidazole resistance in *Tv*, and frequency of reinfection underscore the need for alternative precision therapies^1,6–8^. *Tv* relies on its four anterior and one membrane-bound, recurrent flagellum to propel itself through the genitourinary tract and attach to the mucosa of its human hosts, making the mechanisms driving locomotion potential therapeutic targets^9^. Unfortunately, no high-resolution structures related to *Tv* flagella are currently available, and even tubulin remains uncharacterized in *Tv* despite a putative role in antimicrobial resistance^10–12^.

As observed in low-resolution, thin-section transmission electron microscopy (TEM) studies^13^, the locomotive flagella originate from cytosolic basal bodies, and extend into the flagellar membrane with decorations along the microtubule filaments that stabilize the tubules and facilitate intraflagellar transport. The flagellar core, or axoneme, conforms to the canonical “9+2” axonemal arrangement wherein a central pair of singlet microtubules (MTs) is connected via radial spokes to nine surrounding doublet-microtubules (DMTs) which transduce force through the flagella (Fig. 1)^13,14^. Studies in other organisms revealed DMTs are coated with different combinations of microtubule inner and outer proteins (MIPs and MOPs) that facilitate assembly, stability, and function (Fig. 2A)^15–20^.

**Figure 1.**
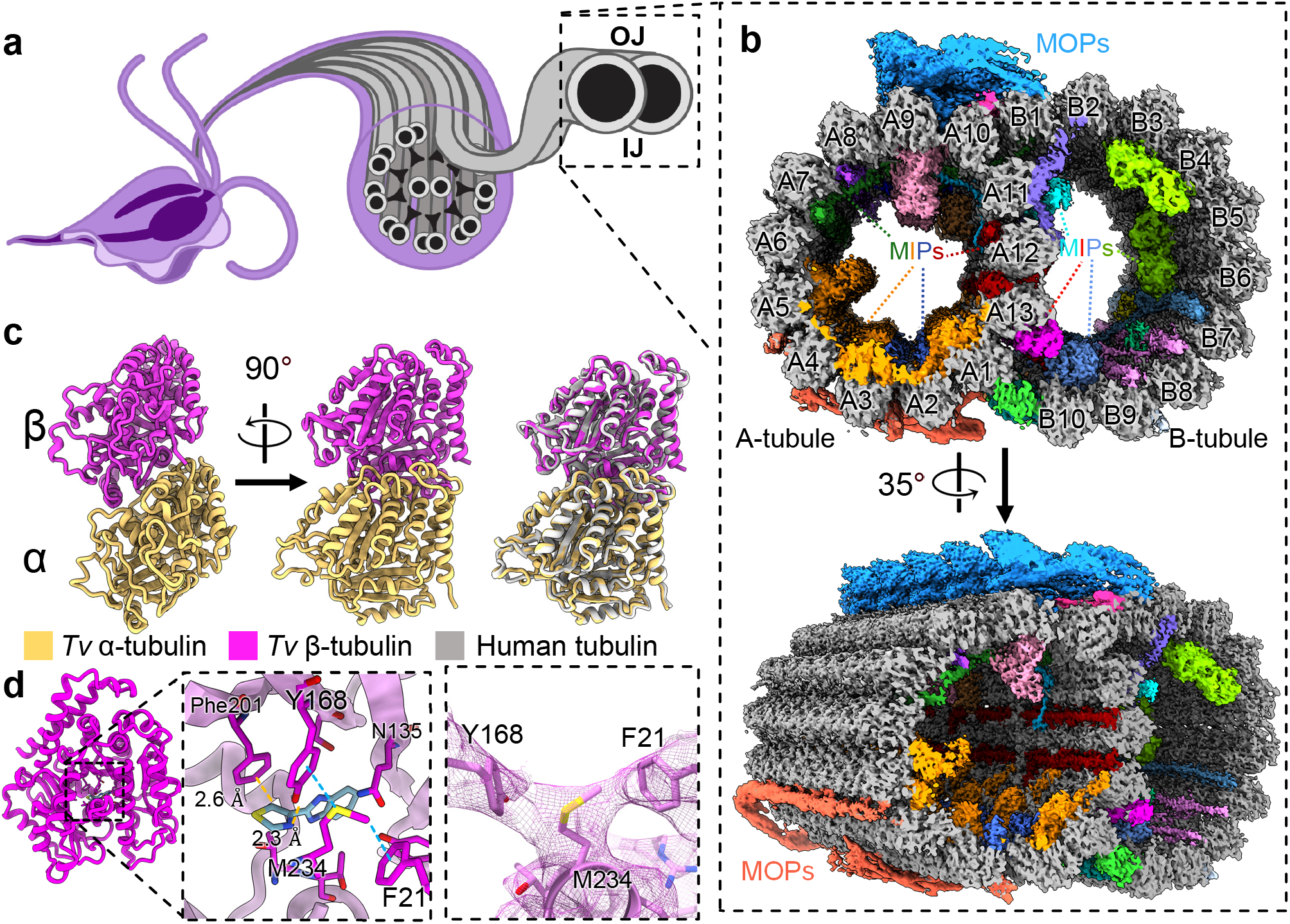
Cryo-EM reconstruction of the doublet microtubules from *Tv*. **(a)** Diagram of axoneme from the flagella of *T. vaginalis.* (**b**) Cross-section of *Tv*DMTs with microtubule inner proteins (MIPs) and microtubule outer proteins (MOPs) indicated with various colors. A- and B-tubules, as well as protofilaments, are labeled. (**c**) Atomic models of α and β tubulin, superimposed with human tubulin (right). **(d)** Alternate view of *Tv* β tubulin (left) and docked thiabendazole molecule (blue) fit into putative binding site with adjacent residues shown (right) and Aro-Met-Aro interaction shown with cryo-EM map density. **IJ:** inner junction; **OJ:** outer junction.

**Figure 2.**
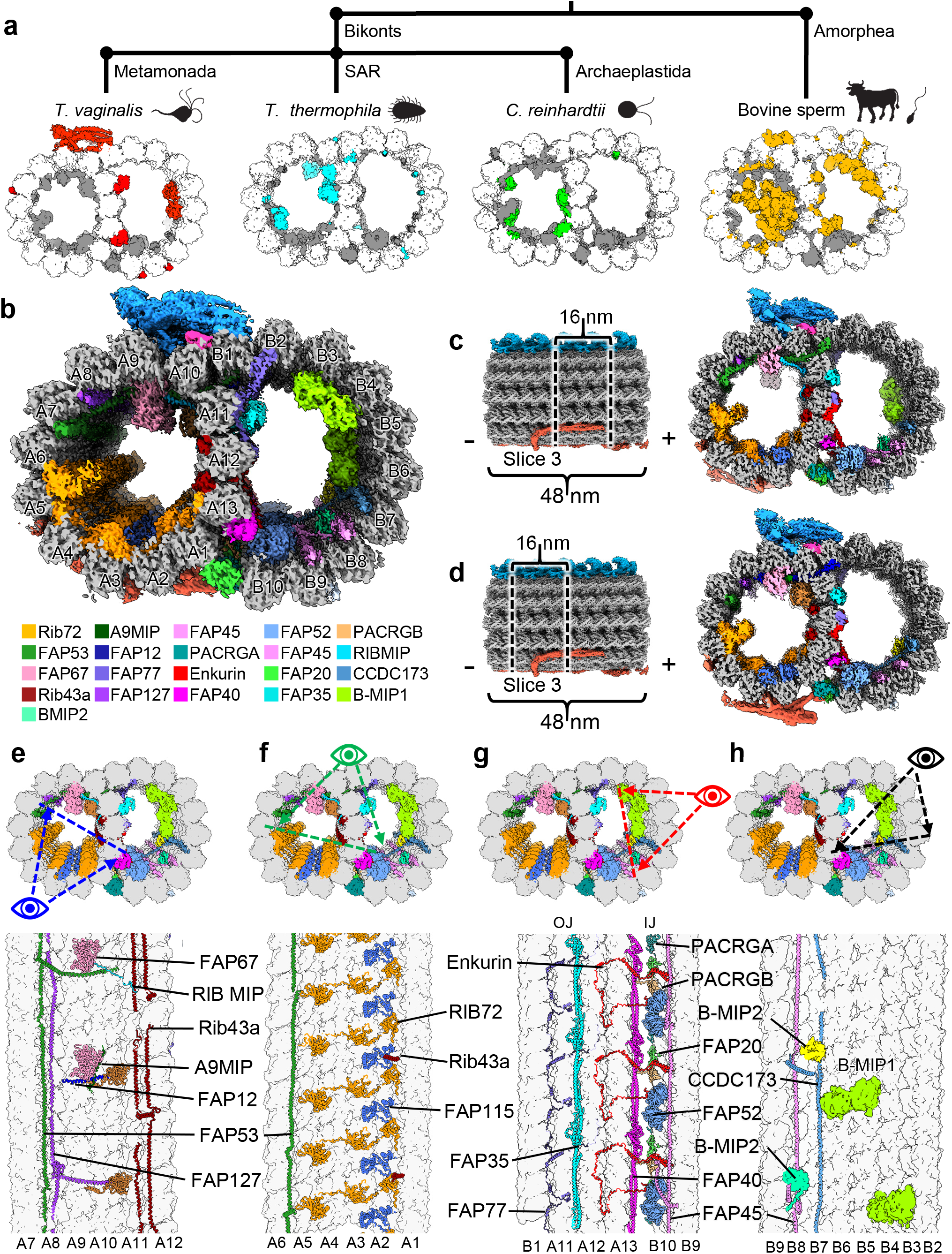
*Tv-*DMTs reveal conserved and novel MIPs. (**a**) Phylogeny tree illustrating proposed divergence between Bikonts and Amorphea (top), with example organisms from these branches and accompanying DMTs (bottom) with tubulin (white), conserved flagella associated proteins (FAPs) (grey), and species-specific FAPs (colored) (**b**) Cross-sectional view of cryo-EM reconstruction of 48 nm repeat with MIP protein densities colored to demonstrate arrangement. (**c and d**) Cross-sectional view of DMTs from the 48 nm repeat map, shown as different 16 nm long sections throughout the DMT. (**e-h**) Cross-sectional views of *Tv*DMTs from different perspectives to illustrate MIP arrangement and periodicity.

Dozens of MIPs and MOPs have been identified across numerous studies of eukaryotic flagella, of which about half are conserved^15–19^. DMTs from multicellular eukaryotes incorporate more complex MIP arrangements, particularly along the highly variable ribbon protofilaments (PFs) that compose the inner and outer junctions (IJ and OJ) where the A- and B-tubules meet^15–19^. In sperm flagella, filamentous tektin bundles near the ribbon PFs are thought to reinforce the long flagella as they swim through the viscous milieu of the genitourinary tract^21,22^. Though the *Tv* genome lacks tektin genes, the parasite swims through the same environment as sperm, coordinating its much shorter flagella into a distinct beating pattern^23^. Despite these apparent differences, it is unclear how the parasite propagates motion under these conditions and suggests a species-specific adaptation which may be exploited for therapeutic development.

Here, we leveraged mass spectrometry, cryogenic electron microscopy (cryo-EM), and artificial intelligence to analyze the DMTs derived from *Tv* parasites and elucidate the structures of the proteins that compose them. Our structure contains 29 atomic models, including the α- and β-tubulin, 18 MIPs and 9 MOPs. Among these, we identified three *Tv*-specific proteins, including one bound to a ligand not observed in the DMTs of other organisms. This first structure from the *Tv* flagella highlights remarkable simplicity in the species’ DMT architecture compared to more complex organisms such as mammals, as well as other protists like *Tetrahymena thermophila*. Despite this simplicity, *Tv* can still traverse the same viscous environment as the more complex mammalian sperm, suggesting a key to parasite locomotion lies in the short list of *Tv-*DMT proteins.

## Results

### *T. vaginalis* DMTs feature both familiar and novel MIPs

We optimized a protocol to isolate DMTs from *T. vaginalis* and limit perturbations to the internal structures, then subjected them to single-particle analysis using cryo-EM. The resultant cryo-EM maps of the 48 nm repeat DMT had a global resolution of 4.2 Å (Fig. 1A) and focused refinement improved local resolution to between 3.2 Å and 3.8 Å (Fig. 1B). Reconstructions of the 16 nm and 96 nm repeat structures were resolved to 3.8 Å and 4.3 Å respectively (Fig. 1A). We also collected mass spectrometry data for our cryo-EM sample to produce a library of potential *Tv*-DMT proteins and utilized cryoID to identify most likely candidates for certain map densities^24^. AlphaFold predicted structures served as initial models for atomic modeling of both conserved and species-specific cryo-EM map densities^25,26^. From our structures we built 29 unique atomic models, including 18 MIPs, 9 MOPs and the α/β tubulin of *Tv* (Supplemental Table 2). Of these proteins, 15 MIPs and all 9 MOPs are conserved between *Tv* and previous DMT structures, whereas 3 MIPs are novel. There are also 5 unassigned MIP and 3 MOP densities that appear to play an important role in DMT function, but for which we lacked sufficient resolution to model.

Consistent with their ∼80% sequence identities, the atomic models of *Tv*’s α- and β- tubulin are nearly identical to those of their human homologs (Fig. 1C), including the region of β- tubulin where many antiparasitic, benzimidazole-derived drugs (BZs) bind (Fig. 1D). Previous studies in *Tv* suggest mutations aromatic residues at codons 168 and 201 in β-tubulin confer BZ resistance^12,27,28^. Indeed, like human β-tubulin’s Phe169 and Tyr202, *Tv* orients Tyr168 and Phe201 into the BZ binding pocket where they are stabilized by Aro-Met-Aro interactions with adjacent Met234 and Phe21 residues and sterically occlude BZ drugs like thiabendazole (TBZ) (Fig. 1D). To corroborate this, we performed docking experiments using AutoDock Vina and found TBZ docked β-tubulin produced large positive binding free energy values (ΔG) (Fig. S2). By contrast a virtual β-tubulin Y168A, P201A mutant exhibited a negative binding free energy when TBZ was docked (Fig. S2). Interestingly, we observe the swapped positions of phenylalanine and tyrosine residues between human and *Tv* β-tubulin, which may help to explain species-specific sensitivity to different BZs.

Like other organisms, the α/β tubulin heterodimers polymerize and assemble into rings of 13 and 10 PFs that compose the A- and B-tubules respectively (Fig. 1B). Within the A-tubule, molecular rulers FAP53, FAP127, and Rib43a impose a 48 nm MIP periodicity and facilitate the organization of other MIPs like FAP67 and RIB72 (Fig. 2E-F). Consistent with studies in *T. thermophila*^18^, FAP115 repeats every 32 nm and creates a mismatch with the 48 nm periodicity of the ruler proteins, leading to 96 nm periodicity (Fig. 2F). Interestingly, FAP141 from other organisms is replaced by the smaller *Tv*FAP12 which lashes FAP67 to the A-tubule lumen like the N-terminal helices of FAP53 and FAP127 (Fig. 2E)^15^. Along with the N-terminal helices of FAP53 and FAP127, *Tv*FAP12 passes into the B-tubule to maintain 16 nm a repeating crosslink between the A- and B-tubules as observed in FAP141 expressing organisms. Unlike other species, the *Tv* ribbon PFs (A11-A13) that divide A- and B-tubules are sparsely decorated with A-tubule MIPs suggesting alternative strategies of ribbon arc stabilization.

In the B-tubule lumen, we found assembly-related MIPs FAP45, CCDC173, enkurin, FAP77, FAP52, FAP20, and PACRGA/B that are conserved amongst other organisms. Interestingly, along the B-tubule side of the ribbon arc, we identified the filamentous MIPs *Tv*FAP35 and *Tv*FAP40, which run lengthwise along the A11 and A13 PFs respectively and may compensate for the dearth of MIPs along the ribbon arc in the A-lumen (Fig. 2G). Further, we observed globular MIPs that span PFs B3-B4 and B5-B6 and exhibit 96 nm periodicity (Fig. 2H). While the map resolution was insufficient to model these proteins, their interactions with the neighboring ruler proteins like CCDC173, indicate an enforced periodicity of 96 nm which is the first of this length from any DMT MIP to date. Though we observed several novel proteins, the *Tv*-DMTs have the simplest MIP organization in the A-tubule with just 10 MIPs (eight identified and two unidentified) compared to the next simplest species of record, *C. reinhardtii*, with 22 A-tubule MIPs^15^. The comparatively simple MIP organization observed in *Tv* suggests the few novel MIPs may play a substantial role in flagellar function.

### *T. vaginalis* microtubules reinforce the inner junction with species-specific protein

The DMT IJs of other organisms are typically composed of FAP52, enkurin/FAP106, PACRG isoforms (PACRGA and PACRGB), and FAP20, while *Tetrahymena* and mammalian DMTs include globular proteins atop FAP52 that mediate interactions with PF A13^18,22,29^. Interestingly, the *Tv*-DMT cryo-EM map revealed the long, filamentous protein *Tv*FAP40, running atop PF A13 at the IJ which alters the topography of this important protofilament. *Tv*FAP40 monomers repeat every 16 nm and are arranged head-to-tail, where head-tail polarity corresponds to the – and +-ends of the DMT respectively (Fig, 3B-C). Each *Tv*FAP40 monomer consists of a globular N-terminal ‘head’-domain (residues 1-145) connected to a coiled-coil ‘tail’ (residues 149-361).

The tail consists of 3 coiled-helices (α7-9) where a proline-rich kink connects α7 and α8 while a 180° turn at the linker between α8 and α9 forms the ‘tip’ of the tail. The tip includes α8 and neighboring residues of α9 (residues 246-282), with both a polar face oriented towards the MT and a hydrophobic face oriented towards a neighboring *Tv*FAP40 monomer (Fig. 3C-D). As the kink reaches into the cleft between tubulin heterodimers, α8 is brought into close contact with tubulin, and establishes electrostatic interactions. The kink also offsets α7 from α8, creating an overhang to bind the head of a neighboring *Tv*FAP40 monomer which may help stabilize the interaction (Fig. 3D).

**Figure 3.**
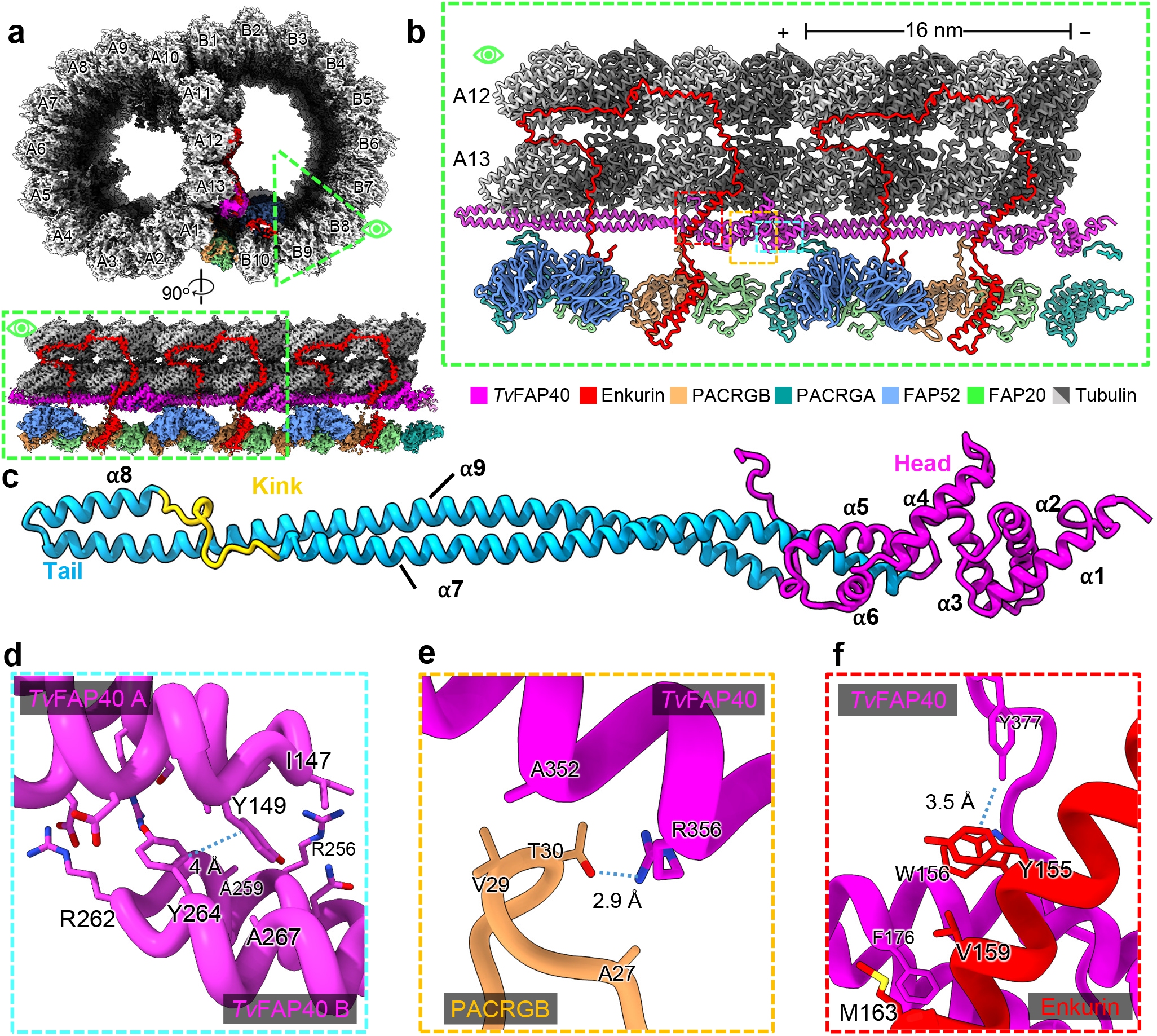
*Tv*FAP40 alters the inner junction arrangement in parasite DMTs. (**a**) Cross-sectional view of cryo-EM reconstruction of 16nnm repeat with protofilaments labeled and proteins near inner junction colored (top) and cutaway view of region of interest (bottom). (**b**) View of atomic models built from map in **a**. (**c**) atomic model of *Tv*FAP40 colored by domain. (**d**) Zoomed-in view of dimerization domain between two *Tv*FAP40 monomers (labeled *Tv*FAP40 A and B). (**e** and **f**) Close-up view of interaction between PACRGB (tan) and *Tv*FAP40 and Enkurin (red), with residues shown to highlight interactions.

*Tv*FAP40’s unique location along PF A13 has not been seen in other MIPs and alters conserved MIP interactions at the inner junction. In other organisms, the PACRGB N-terminus binds the groove between A11 and A13, but in our structure, *Tv*FAP40 blocks this groove and replaces A13 as the binding partner. Additionally, the *Tv*FAP40 C-terminus hooks around α2 of enkurin, where the C-terminal tyrosine (Tyr377) participates in hydrophobic interactions with adjacent aromatic residues from enkurin (Tyr155 and Trp156) (Fig. 3F). This C-terminal hook acts in concert with the *Tv*FAP40 head that binds the other side of enkurin α2 and restricts it such that the bottom end of the helix is 1 nm closer to A13 than in other structures.

### *Tv*-specific FAP40 head domain binds a stabilizing ligand

In addition to binding neighboring monomers, *Tv*FAP40 incorporates a unique ligand binding pocket. Our cryo-EM maps indicate the *Tv*FAP40 head-domain binds a six-pointed, star-shaped ligand, and our atomic model indicates this pocket is positively charged (Fig. 4A-C). Indeed, the putative binding site features seven positively charged side chains oriented towards the points of the star, and density from a likely metal cation coordinated by additional arginine residues (Fig. 4C), which suggests negatively charged functional groups (Fig. 4C & E). Sequence and structural homology searches could not identify similar binding sites, but the high local resolution of our map in this pocket revealed the stereochemistry of the functional groups at the points of the star, consistent with bonding to a non-planar six-membered ring. Together these features suggested an inositol polyphosphate, in this case inositol hexakisphosphate (IP6) which was a good fit for the map density (Fig. 4C).

**Figure 4.**
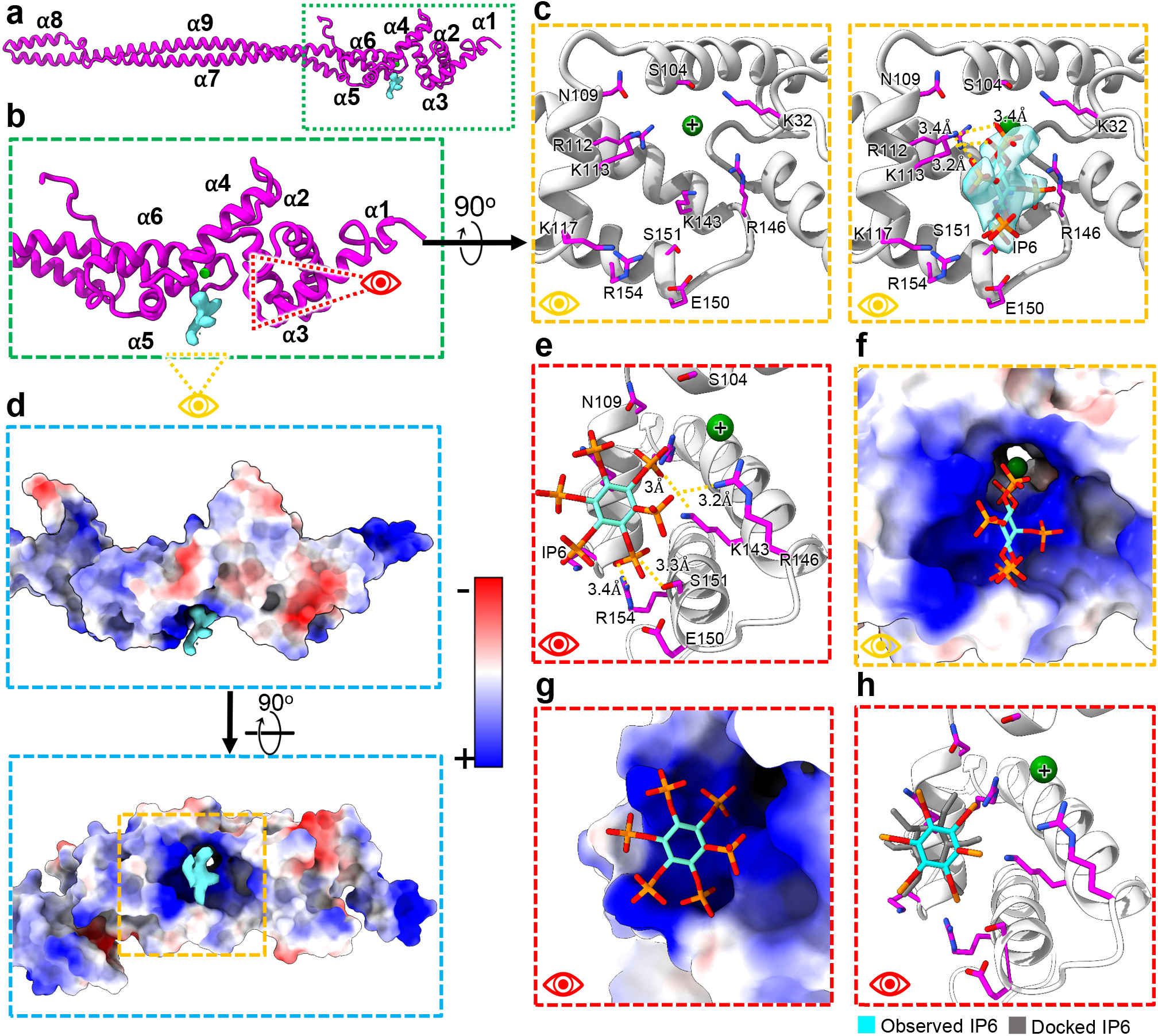
*Tv*FAP40 binds IP6 in a positively charged pocket. (**a**) Atomic model of *Tv*FAP40 with (**b**) zoomed-in view of the head domain and (**c**) perspectives of the putative IP6 binding site with (right) and without (left) IP6 fit into the cryo-EM map. (**d**) Coulombic potential map of head domain from B (top) and rotated (bottom) views with blue and red indicating positive and negative coulombic potentials respectively. (**e**) Side-view of IP6 in binding pocket with adjacent residues shown. (**f-g**) Views from C and E shown with electrostatic potential maps of Tv-FAP40. (**h**) Comparison of observed IP6 binding site and docked IP6.

IP6 is an abundant cellular polyanion known to stabilize positive interfaces such as the pore of HIV nucleocapsids^30^, a trait which may be useful to DMT reinforcing proteins. To confirm whether IP6 was a reasonable ligand assignment, we carried out *in silico* molecular docking using Swissdb’s AutoDock Vina webserver^31–34^. Restricting the docked ligand to the observed binding pocket resulted in docked arrangements consistent with the observed ligand density, and binding energies (ΔG) of -3.5 kcal/mol or less (Fig. 4H). The docking experiments suggested interactions with the same arginine and lysine residues as the real ligand structure in the binding pocket. These results support the notion that IP6 acts as a ligand within *Tv*FAP40 and may stabilize the head to reinforce its interactions with enkurin and the tail of its neighboring monomer and microtubule PF. Interestingly, in zebrafish embryos the IP6 producing enzyme (Ipk1) was found to localize to basal bodies of cells, and Ipk1 knockdown disrupted cilia growth and beating^35,36^. Combined with our structures, these studies point to an uncharacterized role for IP6 in flagellar function and stability.

### *Tv*FAP35 secures FAP77 and buttresses the outer junction of *Tv*-DMTs

Directly above *Tv*FAP40 in the B-tubule we identified a novel, filamentous density along the ribbon arc PF A11 as *Tv*FAP35, another *Tv*-specific protein (Fig. 5A-C). In the B-tubule, *Tv*FAP35 repeats every 16 nm in a head-to-tail fashion with the heads and tails oriented to the – and +-ends respectively (Fig. 5B). The *Tv*FAP35 tail domain has the same ‘kinked-coiled-coil’ fold as *Tv*FAP40 including the tips that mediate MT binding and dimerization (Fig. 5C-F). The head-domain of *Tv*FAP35 differs from *Tv*FAP40, as it includes only a flexible N-terminus and helix-turn-helix (Fig. 5C), as opposed to the six helices found in the *Tv*FAP40 head (Fig. 3C).

**Figure 5.**
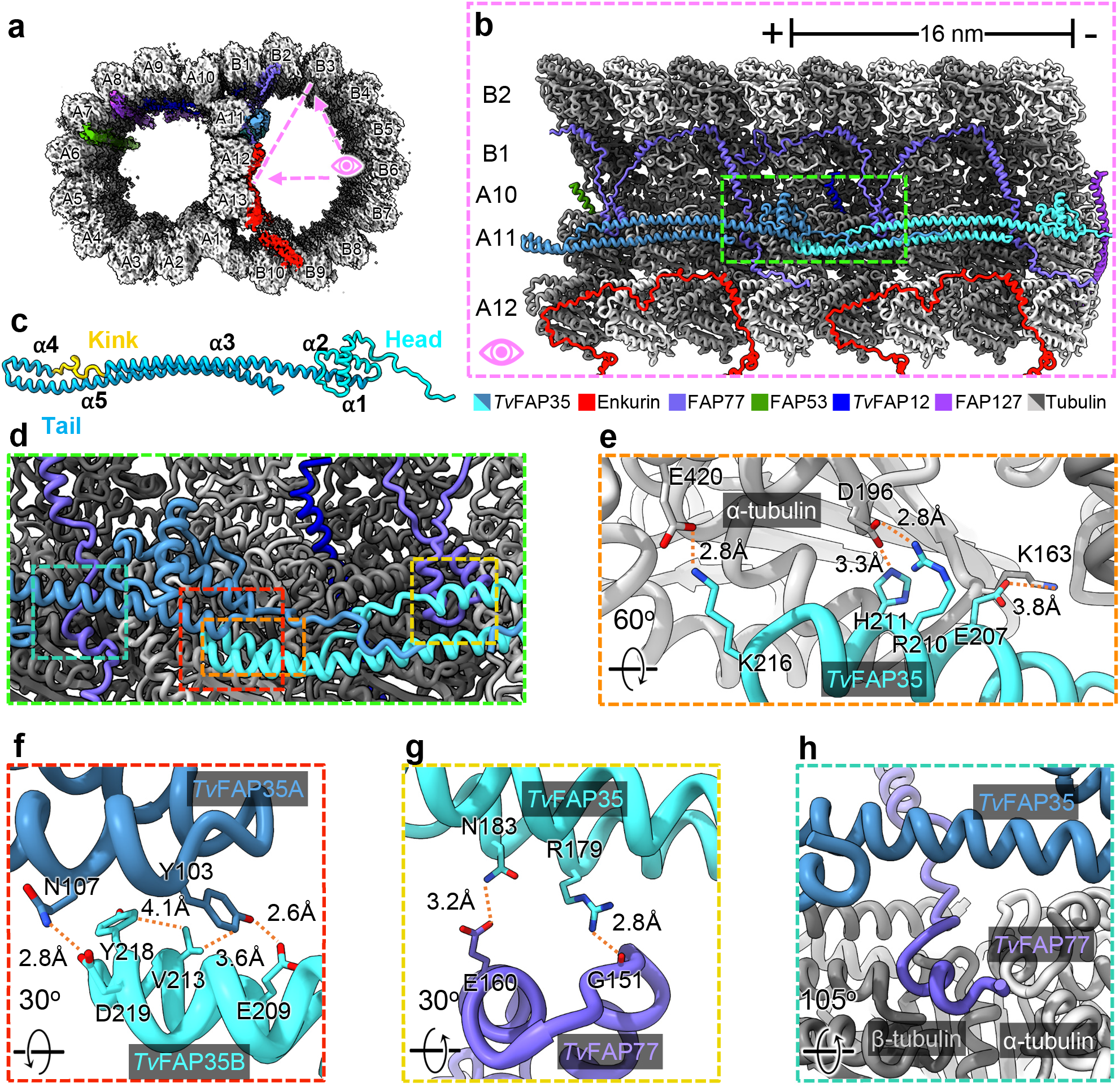
*Tv*FAP35 stabilizes ribbon PF A11 and outer junction proteins. (**a**) Cross-sectional view of the *Tv*DMT cryo-EM map with enkurin and outer junction proteins colored. (**b**) 32 nm section of protofilaments A10, A11, A12, B1, and B2, along with their associated MIPs, shown with atomic models. (**c**) *Tv*FAP35 monomer labeled with head (cyan), tail (blue), and kink (yellow), with helix numbers. (**d**) Zoomed-in view including important interactions of *Tv*FAP35. (**e**) Electrostatic interactions at the MT-binding motif of *Tv*FAP35. (**f**) Mixed residue interactions at the dimerization interface between *Tv*FAP35 monomers. (**g**) Interactions between TvFAP35 and the helix-turn-helix (residues 140-164) of *Tv*FAP77. (**h**) Residues 238-246 of *Tv*FAP77 pass near the *Tv*FAP35 coiled-coil. Residues 255 and after of *Tv*FAP77, which stretch further down, are omitted for clarity.

The position of *Tv*FAP35 along A11 is similar to that of tektin-like protein 1 (TEKTL1) which is thought to reinforce OJ stability during flagellar beating in sperm DMTs^22^. Further, the coiled-coil structure of *Tv*FAP35 resembles the 3-helix bundle architecture of TEKTL1. Unlike TEKTL1, the coiled coils of *Tv*FAP35 include a proline-rich kink that occupies the cleft between tubulin heterodimers. As PFs bend, gaps form at the interface between tubulin heterodimers^37^, and the *Tv*FAP35 kink may create stress relief points along A11 by acting as a flexible linker which accommodates bending. Thus, like TEKTL1, the coiled-coils of *Tv*FAP35 may provide structural stability to the DMT while the kink allows bending and greater flexibility. *Tv*FAP35 also interacts with FAP77, a MIP that aids in B-tubule formation and tethers complete A- and B-tubules together at the OJ^18^. The FAP77 helix-turn-helix motif (residues 140-164) is braced to PF A11 via electrostatic interactions with the coiled-coil of *Tv*FAP35 (Fig. 5B, C, F). Further, the tail-domain of *Tv*FAP35 passes over residues 238-246 of *Tv*FAP77 which run between a cleft of the A11 PF and reinforces *Tv*FAP77’s association to A11 (Fig. 5G). These observations suggest that, like *Tv*FAP40, *Tv*FAP35 plays an integral role in the stabilization of the ribbon PFs and their associated MIPs. Additionally, because FAP77 is implicated in B-tubule assembly^18^, the interactions of *Tv*FAP35 with FAP77 and A11 suggest that *Tv*FAP35 may also contribute to DMT assembly.

### Novel *Tv* proteins share an ancient MT binding motif

Upon comparison, we noticed both *Tv*FAP35 and *Tv*FAP40 have kinked-coiled-coils composed of three helices (ɑ1-3, Fig. S2) with similar lengths, dimerization domains, and MT binding motifs (Fig. 3-4). This similarity prompted us to search for homologous proteins via amino acid sequence alignment, but this returned few candidates. Interestingly, the coiled-coils of *Tv*FAP35 and *Tv*FAP40 share just 23% identity despite similar folds. We next turned to structural alignment using *FoldSeek*^38^, and identified numerous structural homologs. After curating homolog candidates by removing those without kinked-coiled-coil domains or with TM-scores below 0.4, 31 homolog candidates were selected for further comparison. Initial analysis revealed all kinked-coiled-coil containing homologs belonged to the group *<u>Bikonta</u>*, which includes many protists, and excludes animals, fungi and amoebozoans.

Six kinked-coiled-coil homologs from protists, representing different clades were selected for multiple sequence alignment with *Tv*FAP35 and *Tv*FAP40 (Fig. S3A) which revealed several conserved residues from the dimerization and MT binding domains. Based on the *Tv*FAP35 sequence, the dimerization domains include hydrophobic residues at Val213 and aromatic residues at Tyr103 and Tyr218, which form hydrophobic interfaces between neighboring monomers (Figs. S2G). Further, the MT binding motif on α2 has a high proportion of charged residues, which are likely important in tubulin binding (Fig. S3F). Outside of the dimerization and MT binding motifs, the kinked-coiled-coils exhibit an average sequence conservation of ∼20% which may be necessary to accommodate different MIPs, as observed in our novel proteins (Fig. 3 and 5).

### *T. vaginalis* microtubule outer proteins exhibit 8 nm periodicities

Considering the marked simplicity of the *Tv*-DMT MIP arrangement, we expected to find comparably simple MOP organization. Along the A-tubule we observed both the canonical N-DRC and radial spoke complexes that mediate inter-axoneme connections and flagellar bending (Fig. 6A-C). We see that, like other DMT structures, the axoneme-related proteins exhibit 96 nm periodicity enforced by the molecular ruler proteins CCDC39 and CCDC40, which coil their way between PFs A3 and A2 (Fig 6B-E). Besides N-DRC and radial spoke proteins, a diverse arrangement of filamentous MOPs occupies the clefts and the surface of several PFs. Previous DMT structures from other species found the shortest MOP periodicity to be 24 nm^18,22^.

**Figure 6.**
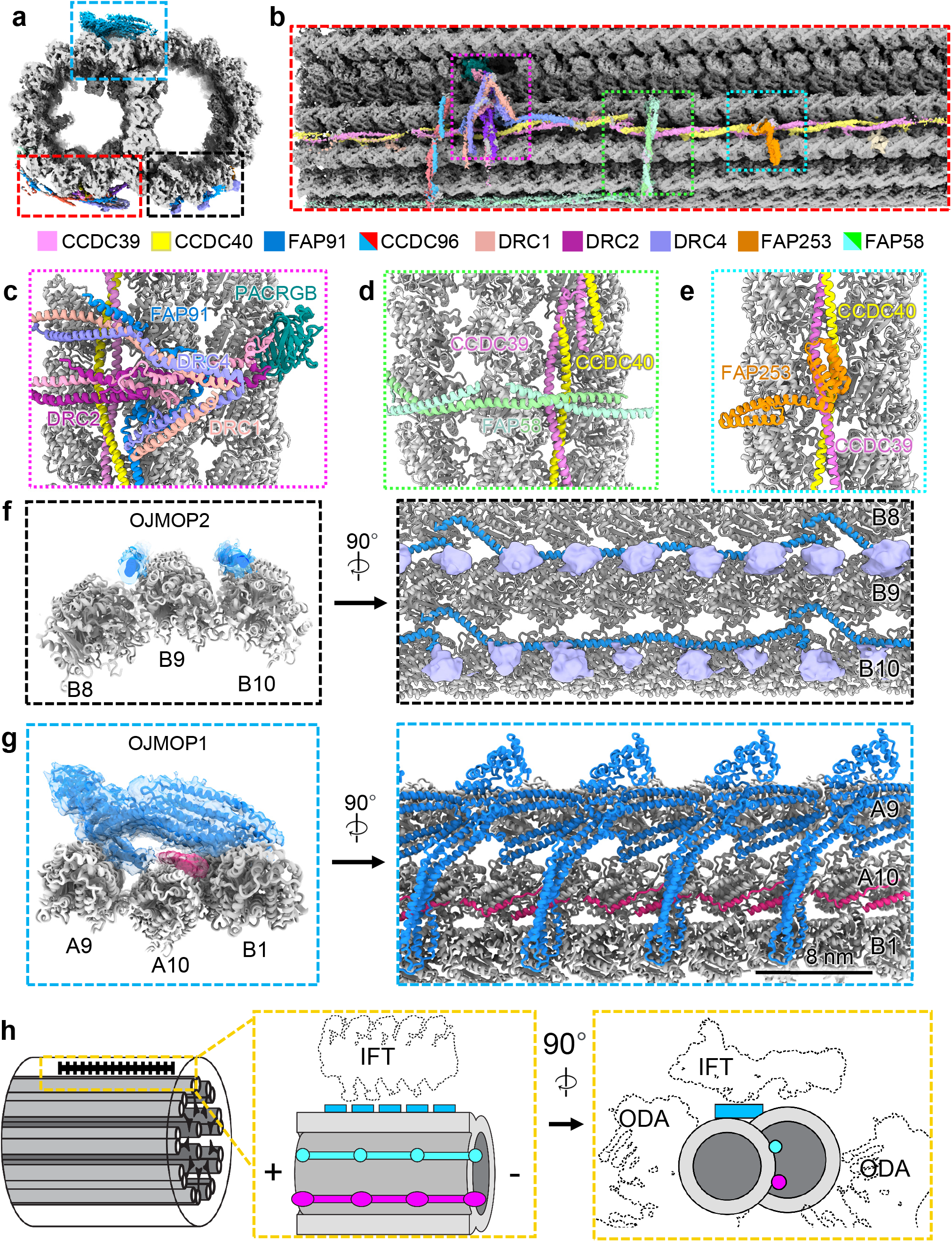
Microtubule organization reveals novel 8nm periodicity. (**a**) Cross-sectional view of 96 nm repeat map, colored by MOP. (**b**) external view of *Tv*DMT and zoomed in views of MOPs (**c-e**). (**g**) *Tv*OJMOP2 demonstrating 8nm periodicity with cross-sectional (left) and external views (right). (**f**) *Tv*OJMOP1 demonstrating 24 nm periodicity as cross-section (left) and external view (right). (**h**) Schematic view of *Tv*-DMT organization with dotted lines to indicate positions of IFT and inner and outer dynein arm attachment (IDA and ODA).

*Tv*MOP1 is 24 nm repeating MOP that arranges head-to-tail in the furrows between PFs A3, A4, B8, B9, & B10 and contacts the flexible C-terminal tails of α and β tubulin in the B-tubule (Fig. 6F). Interestingly, though the exterior of the outer junction is sparsely decorated in DMTs structures from other organisms^15,22^, we found this area to contain a large filamentous protein that repeats every 8 nm and a smaller filament that runs in a zig-zag beneath it and between A10 and B1 (Fig. 6F). The large protein density fashions an ankyrin-like domain seated atop a large coiled-coil domain, which spans the gap between PF A9 and B1 (Fig. 6F).

Due to limited local resolution, we were unable to confidently assign the identities of these proteins and instead dubbed them *Tv* outer junction microtubule outer protein 1 and 2 (*Tv*OJMOP1 and *Tv*OJMOP2) for the large and zig-zag MOPs respectively. *Tv*OJMOP1 exhibits an 8 nm periodicity like that of tubulin heterodimers, an unusual repeat length amongst DMT MOPs that crosslinks PF B1 to A9 and A10 (Fig. 6G). *Tv*OJMOP1 was observed in only 1/5 of particles, suggesting that some may have been lost during DMT isolation or that *Tv*OJMOP1 localizes to certain regions of the axoneme. Exhaustive search through AlphaFold predicted structures from our proteomic data using a strategy similar to that of DomainFit^40^ yielded the following 5 candidate proteins which contain both ankyrin and coiled-coil domains: TVAGG3_0305310, TVAGG3_0421180, TVAGG3_0431750, TVAGG3_0596110, and TVAGG3_0415080. However, none of these candidates could fully account for the observed density, and so it remains unclear if *Tv*OJMOP1 is composed of one or more of these proteins. Recent work in *C. reinhardtii* has demonstrated that anterograde intraflagellar transport (IFT) brings IFT-B complexes directly over this area (Fig. 6H)^41^. However, as components are often lost during DMT isolation they are unlikely candidates. As it features an ankyrin domain oriented towards the would-be IFT-B cargo (Fig. 6G-H), *Tv*OJMOP1 may interact with TPR-rich proteins of IFT-B to stabilize the cargo. Additionally, others have documented the tendency for cytoplasmic dynein motors to jump between PFs^42^. *Tv*OJMOP1 may therefore create tracks to keep the dynein motors on their preferred A-tubule PFs.

## Discussion

*T. vaginalis* pathogenesis relies on the parasites’ locomotive flagella to establish infection and spread between human hosts^23,43^. This study reports the first high-resolution structure of *Tv* flagellar doublet microtubules, elucidating their molecular composition, architectural arrangement, atomic structures, and small molecule ligands. In addition to the first atomic structures of the *Tv* tubulin subunits comprising the DMTs, we have identified 20 MIPs and 13 MOPs distributed across the A- and B-tubules. These MIPs and MOPs mediate *Tv*-DMT function in the flagella with several novel proteins. As the first near-atomic structure of flagellar microtubules in the major human parasite *Trichomonas vaginalis*, our results provide a structural framework to understand the parasite’s distinct locomotion, offer insights into anthelmintic drug resistance, and identify new targets for precision medicine.

With a relatively short list of both conserved and *Tv*-specific MIPs, the *Tv*-DMT is perhaps the simplest among known DMT structures. Notably, the *Tv* A-tubule fashions the fewest MIPs of any characterized organism (Fig. 2). Among them, the ruler proteins Rib43a, FAP53, and FAP127 are conserved, but lack many of the interacting partners of their homologs in other species, such as mammalian sperm, that traverse the same environment. The sparsity of A-tubule ribbon proteins in *Tv* suggests these proteins are less essential for locomotion in the human genitourinary tract, which contrasts with the complex MIP arrangement of sperm-specific proteins and tektin bundles seen in the A-tubule of mammalian sperm^22,29^. While *Tv* and sperm exhibit distinct flagellar beating patterns, the sinusoidal beating pattern of the recurrent flagella in *Tv* suggests the additional MIP complexity observed in sperm is not essential to this style of beating. However, human sperm swim five times faster than *Tv* and must propagate beating over longer flagella, so sperms’ complex A-tubule MIP arrangement may facilitate rapid propulsion through their viscous environment^9,44^.

Remarkable specialization is observed in the B-tubule, where several novel proteins reinforce the ribbon arc in a manner similar to tektin bundles from other organisms^15^. Like tektin, the *Tv*FAP35 and *Tv*FAP40 proteins exhibit 16 nm periodicity and similarly interact with other MIPs along their respective protofilaments (Fig. 3 and 5). However, unlike tektin, *Tv*FAP35 and *Tv*FAP40 have variable head domains which seemingly confer different functionalities. To this end, the positively charged pocket of *Tv*FAP40 putatively binds an IP6 pocket factor (Fig. 4). In HIV, IP6 acts as a pocket factor to stabilize the nucleocapsid lattice^30^. Considering *Tv*FAP40 likely plays a stabilizing role along the ribbon arc, IP6 binding may augment that stabilization by reinforcing the interactions between the monomers at the head-tail interface. Binding abundant biomolecules is a common strategy amongst pathogens, particularly viruses^45–47^, but this is the first instance such pocket factors have been documented in DMTs.

The *Tv*-specific MIPs and MOPs are particularly significant in light of their role in propagating the pathogenesis of the most widespread non-viral STI^3^. Specialization differentiates the parasite’s DMTs from those of other organisms, including their human hosts, thus drugs targeting these specialized components would have minimal toxicity. For instance, the unique cofactor binding pocket found in the *Tv*-specific *Tv*FAP40 protein has a structure with no known homologs and appears to specifically bind IP6 (Fig. 4). This pocket could be targeted by antimicrobial compounds to destabilize parasite DMTs with limited off-target effects. Notably, the only homologous proteins to *Tv*FAP40 belonged to other Bikonts, and include other human-borne parasites like *T. brucei* and *Leishmania donovani*, that may incorporate similar species-specific proteins (Fig. S3). While this study represents the first of its kind on the DMT from a human-borne parasite, it has demonstrated the power of *in situ* cryo-EM over other structural or *in silico* methods, to open new avenues for rational drug design. Together, our findings provide a basis to explore the contribution of microtubule-associated proteins to the unique aspects that allow *T. vaginalis* to swim through the human genitourinary tract, and the diversity of eukaryotic motility in general. Moreover, the atomic details revealed in species-specific proteins and bound small molecules can inform the rational design of therapeutics.

## Author contributions

Z.H.Z and P.J. designed and supervised the project. K.A.M., E.W., and A.S. prepared samples. E.W. conducted mass spectrometry work. A.S. and S.K. performed cryo-EM imaging and prepared 3D reconstructions. Under the guidance of Z.H.Z., A.S., S.K., and E.C. built the atomic models, interpreted the structures, and made the figures and wrote the paper; all authors reviewed and approved the paper.

## Acknowledgements

This work was supported in part by grants from the National Institutes of Health (R01GM071940 to Z.H.Z. and R01AI103182/R33AI119721 to P.J.J.). K.A.M. and A.S. received support from NIH Ruth L. Kirschtein National Research Service Award AI007323. We acknowledge the use of resources at the Electron Imaging Center for Nanomachines supported by UCLA and by instrumentation grants from NIH (1S10RR23057, 1S10OD018111) and NSF (DBI-1338135 and DMR-1548924). We acknowledge support from the UCLA AIDS Institute, the James B. Pendleton Charitable Trust, and the McCarthy Family Foundation.

## Methods and Data Availability

### Cell culture

*T. vaginalis* strain G3 was cultured in Diamond’s modified trypticase-yeast extract-maltose (TYM) medium supplemented with 10% horse serum (Sigma-Aldrich), 10 U/mL penicillin, 10 µg/ml streptomycin (Gibco), 180 µM ferrous ammonium sulfate, and 28 µM sulfosalicylic acid^48^. 2L of parasites, grown at 37 °C and passaged daily, were harvested by centrifugation, and washed twice with phosphate-buffered saline and pelleted at low speed. Cells were resuspended in 50 mL lysis buffer (2% IGEPAL CA-630, 2% Triton X-100, 10% glycerol, 10 mM Tris, 2 mM EDTA, 150 mM KCl, 2 mM MgSO4, 1 mM dithiothreitol [DTT], 1× Halt protease inhibitors [pH 7.4]) and lysed in a Stansted cell disrupter, operated at 30 lb/in^2^ front pressure and 12 lb/in^2^ back pressure.

Cytoskeletal elements were harvested similar to what has been previously described^49^. Lysates were recovered and maintained at 4 °C for all subsequent steps. Nuclei were removed via low-speed centrifugation (1000 x g) for 10 mins to generate pellet 1 (P1) and lysate 1 (L1). L1 was centrifuged (10,000 x g for 40 mins) to pellet cytoskeletal components into P2 and L2. Cytoskeleton pellets (P2) were resuspended in 1 mL low salt (LS) buffer (150 mM NaCl, 50 mM Tris, 2 mM MgCl2, 1 mM DTT, 1× complete protease inhibitor (Sigma-Aldrich)) and centrifuged at low speed (1000 x g, 10 mins) to pellet cellular debris into P3. The resulting lysate (L3) was placed a sucrose cushion (30% w/v sucrose in LS buffer) and centrifuged at low speed (1,800 x g, 10 mins). The supernatant atop the cushion was collected and resuspended in 1 mL LS buffer prior to centrifugation (5,000 x g 15 mins) to pellet larger cytoskeletal components (P4). The lysate was finally centrifuged at high speed (16,600 x g, 40 minutes) to pellet axoneme related cytoskeletal elements (P5). The P5 was then resuspended in minimal volume of LS buffer supplemented with 5 mM ATP and left at RT for 1 hour.

### In-solution digestion, Mass Spectrometry Data Acquisition and Analysis

*T. vaginalis* cytoskeleton pellets P4 and P5 resuspended in low salt (LS) buffer were mixed with 4× volume of ice-cold acetone and kept at -20°C for 2 h. The mixtures were centrifuged at 4°C with 14,000 rpm for 15 min and supernatants discarded. The air-dried pellets were fully dissolved in 8 M Urea in 100 mM Tris-HCl (pH 8) at 56 °C and the proteins reduced with 10 mM Tris(2-carboxyethyl) Phosphine for 1 h at 56 °C. The reduced proteins were then alkylated with 40 mM iodoacetamide for 30 min in dark at room temperature and the reaction was quenched with Dithiothreitol at a final concentration of 10 mM. The alkylated samples were subsequently diluted with 7× volume of 100 mM Tris-HCl pH 8, to 1M Urea concentration. To generate peptides, Pierce Trypsin Protease (Thermo Fisher Scientific) was added to the samples and the ratio of trypsin:protein was 1:20 (w/w). The digestion reaction was incubated at 37 °C overnight, and the residue detergents in the protein samples were removed using a HiPPR Detergent Removal Spin Column Kit (Thermo Fisher Scientific) on the next day. Prior to the mass spectrometry assay, the samples were desalted with Pierce C18 Spin Columns (Thermo Fisher Scientific) and lyophilized.

Three biological replicates were prepared and trypsin-digested following the steps above for fractions P4 and P5, respectively. The lyophilized protein pellets were dissolved in sample buffer (3% Acetonitrile with 0.1% formic acid) and ∼1.0 µg protein from each sample was injected to an ultimate 3000 nano LC, which was equipped with a 75µm x 2 cm trap column packed with C18 3µm bulk resins (Acclaim PepMap 100, Thermo Fisher Scientific) and a 75µm x 15 cm analytical column with C18 2µm resins (Acclaim PepMap RSLC, Thermo Fisher Scientific). The nanoLC gradient was 3−35% solvent B (A = H2O with 0.1% formic acid; B = acetonitrile with 0.1% formic acid) over 40 min and from 35% to 85% solvent B in 5 min at flow rate 300 nL/min. The nannoLC was coupled with a Q Exactive Plus orbitrap mass spectrometer (Thermo Fisher Scientific, San Jose, CA), operated with Data Dependent Acquisition mode (DDA) with inclusion list for the target peptides. The ESI voltage was set at 1.9 kV, and the capillary temperature was set at 275 °C. Full spectra (m/z 350 - 2000) were acquired in profile mode with resolution 70,000 at m/z 200 with an automated gain control (AGC) target of 3 × 106. The most abundance 15 ions were subjected to fragmentation by higher-energy collisional dissociation (HCD) with normalized collisional energy of 25. MS/MS spectra were acquired in centroid mode with resolution 17,500 at m/z 200. The AGC target for fragment ions is set at 2 × 10^4^ with maximum injection time of 50 ms. Charge states 1, 7, 8, and unassigned were excluded from tandem MS experiments. Dynamic exclusion was set at 45.0 s.

The raw data was searched against total *T. vaginalis* annotated proteins (version 63) downloaded from TrichDB, using ProteomeDiscoverer 2.5. Following parameters were set: precursor mass tolerance ±10 ppm, fragment mass tolerance ±0.02 Th for HCD, up to two miscleavages by semi trypsin, methionine oxidation as variable modification, and cysteine carbamidomethylation as static modification. Protein abundance was quantified using Top 3 approach, i.e., the sum of the three most intense peptides coming from the same protein. Only proteins that were detected in all three replicates of P4 or P5 were included for further analyses, which resulted in a total of 386 and 311 proteins identified from P4 and P5, respectively. Among these common proteins, contaminants that are obviously not cytoskeletal proteins were identified from the datasets based on the GO terms and function annotations. For instance, proteins annotated as histone, kinase or DNA binding proteins or proteins located in subcellular compartments e.g., translational apparatus, nucleus, plasma membrane, were removed from the datasets. As a consequence, the numbers of putative cytoskeletal proteins identified in P4 and P5 were reduced to 303 and 239, respectively. The union of dataset P4 and P5, which consists of 371 distinct proteins, represent the entire cytoskeletal proteome of *T. vaginalis* identified by this study. DeepCoil 2.0 program was employed to predict coiled-coil domains (ccds) from the 371 putative cytoskeletal proteins based on protein sequence^50^. Three indices, i.e., number of ccds within each protein, the average length of ccds in each protein and percentage of total protein length occupied by ccds were calculated based on the output of DeepCoil 2.0. In addition to the cytoskeletal proteome in this study, the presence of ccds was also investigated for the hydrogenosome proteome of *T. vaginalis* and a randomly picked *T. vaginalis* protein dataset ^51^.

### Cryo-EM sample preparation and image aquisition

To prepare DMTs for single particle analysis, 2.5 µL of DMT lysate was applied to glow discharged carbon holey grids (R2/1) (Ted Pella) and incubated on the grid for 1 minute prior to blotting and plunge freezing into a 50:50 mixture of liquid ethane and propane using a Vitrobot Mark IV (Thermo-Fisher). Flash frozen grids were stored under liquid nitrogen until cryo-EM imaging.

Dose fractionated cryo-EM movies were recorded on a K3 direct electron detector (Gatan) equipped Titan Krios electron microscope (FEI/Thermo-Fisher) fitted with a Gatan Imaging Filter (GIF) and operated at 300 keV. Movies were recorded at a nominal magnification of 81,000 x and calibrated pixel size of 0.55 Å at the specimen level, operated in super resolution mode. Using SerialEM^52^, 30,834 movies were recorded with a cumulative electron dose of ∼ 45 e^-^/A^2^.

### Cryo-EM image processing and 3-dimensional reconstruction

Movie frame alignment and motion correction were performed in CryoSPARC^53^, to generate cryo-EM micrographs from each movie. Patch-aligned and dose weighted micrographs were binned 2X to improve processing speeds and transferred for processing in Relion 4.0 and Topaz automated particle picking, using the filament option “-f” incorporated by Scheres and colleagues^54–56^. Picked particles coordinates were extracted in Relion using the particle extract job with helical option enabled to extract particles every 8.2 nm along the picked filaments. The extracted particles were transferred back to CryoSPARC for further analysis and 3D reconstruction. 942,986 DMT particles were initially screened for quality using 2D classification job type, and those classes with good features were chosen for further data processing leaving 868,683 particles. Initial 3D reconstructions were made using 2X binned particles to expedite data processing. CryoSPARC’s Helix refine job type was used to refine the DMT particles and prevent particles from the same filament from being placed in different half-sets during refinment. With half sets determined, the particles were then subjected to non-uniform refinement to yield an initial DMT reconstruction based on the 8.2 nm repeating tubulin heterodimer organization.

We next carried out focused classification and refinements as described previously^15^, using CryoSPARC. Briefly, cylindrical masks over MIPS or MOPs with known periodicities were used to relax the 16, 48, and 96 nm periodicity from the initial 8.2 nm repeating DMT structure in stepwise fashion. To improve local resolutions, we performed focused local refinements wherein cylindrical masks were placed over specific protofilaments so that CryoSPARC could be used to align those protofilaments and their MIP and MOP features. This resulted in 8, 16, 48, and 96 nm reconstructions with 3.8, 3.8, 4.2, and 4.3 Å global resolutions, respectively. Local resolutions were improved using the local refinement job types in CryoSPARC, with maps over the regions of interest.

### Atomic Modeling and Docking

The tubulin models were built using AlphaFold predicted models of α- and β-tubulin and using molecular dynamics flexible fitting software in UCSF ChimeraX^57,58^. To model MIPs, homologs from other organisms with existing structures roughly fit into our DMT maps before using NCBI’s basic local alignment search tool (BLAST) to identify homologs in *Tv* and confirmed their identity using our mass spectrometry data^59^. For densities lacking homologous proteins, initial models were built using DeepTracer^60^, followed by refinement in Coot^61^.

The identities of unknown densities were confirmed using automated building in ModelAngelo and standard Protein BLAST of the predicted amino acid sequences against TrichDB database^59,62,63^. Alternatively, or often in combination with ModelAngelo predicted models, cryoID was used to identify the most likely candidates for cryo-EM densities^24^. Further attempts to fit proteins in low resolution regions were made using a strategy similar to that of the DomainFit software package^40^. Briefly, visual inspection of AlphaFold predicted structures also aided in matching of candidates with map density shapes to assess potential matches. Models were fit using Coot and ISOLDE as described previously ^57,64^ and refined using Phenix Real Space Refinement^65^.

Docking of thiabendazole (SMILES: C1=CC=C2C(=C1)NC(=N2)C3=CSC=N3) into *Tv* β- tubulin was performed using SwissDock tools and AutoDock Vina version 1.2.0^32,33^. Tyr168 and Phe201 were mutated to alanine residues using the “swap amino acid” function in UCSF ChimeraX^58^. The box center was placed at 361 – 472 – 277 for each run with dimensions 10 – 10 – 15 and sampling exhaustivity set to the default value of 4.

We used the same software as above for docking IP6 (SMILES: C1(C(C(C(C(C1OP(=O)(O)O)OP(=O)(O)O)OP(=O)(O)O)OP(=O)(O)O)OP(=O)(O)O)OP(=O)(O)O) into *Tv*FAP40, except that the box center was placed at 393 – 277 – 260 and box size was left at default of 20 – 20 – 20 with sampling exhaustivity of 4. Grid box size was chosen to constrain ligands to putative binding sites from previous studies (thiabedazole) or observed localization (IP6) ^27^. Structure visualization and figure preparation were done with UCSF ChimeraX^58^ and Adobe illustrator, respectively.

### Data availability

Cryo-EM maps of the 16, 48, and 96 nm repeats have been submitted to the Electron Microscopy Data Bank and can be found under accession numbers EMD-XXXXX, EMD- XXXXX, and EMD-XXXXX respectively. The coordinates for the complete atomic models were deposited in the Protein Data bank under accession number XXXX.

---

**Figure S1.**
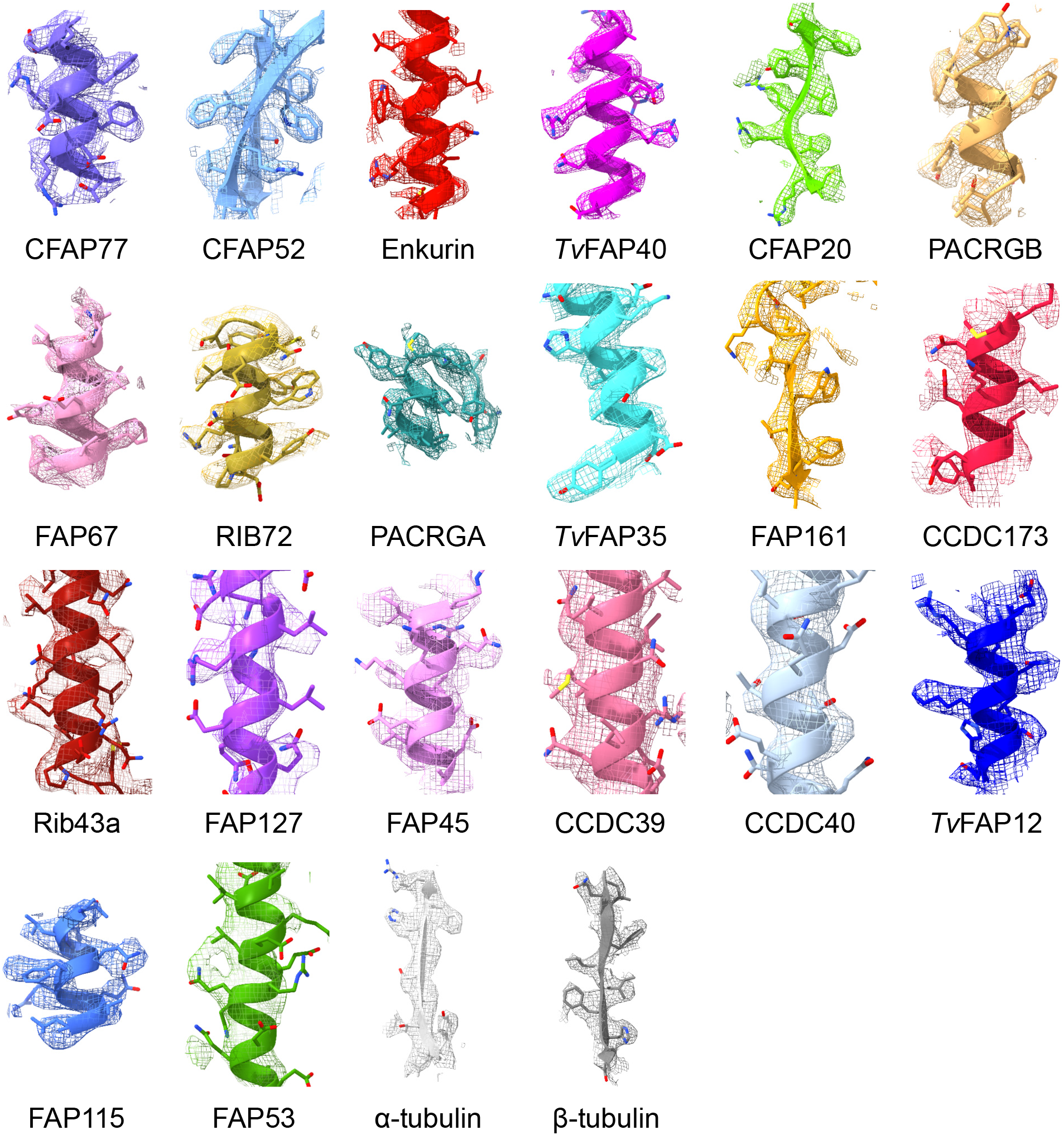
Fitted models in cryo-EM densities. Examples of cryo-EM maps with fitted atomic models of MIP and MOP proteins.

**Figure S2.**
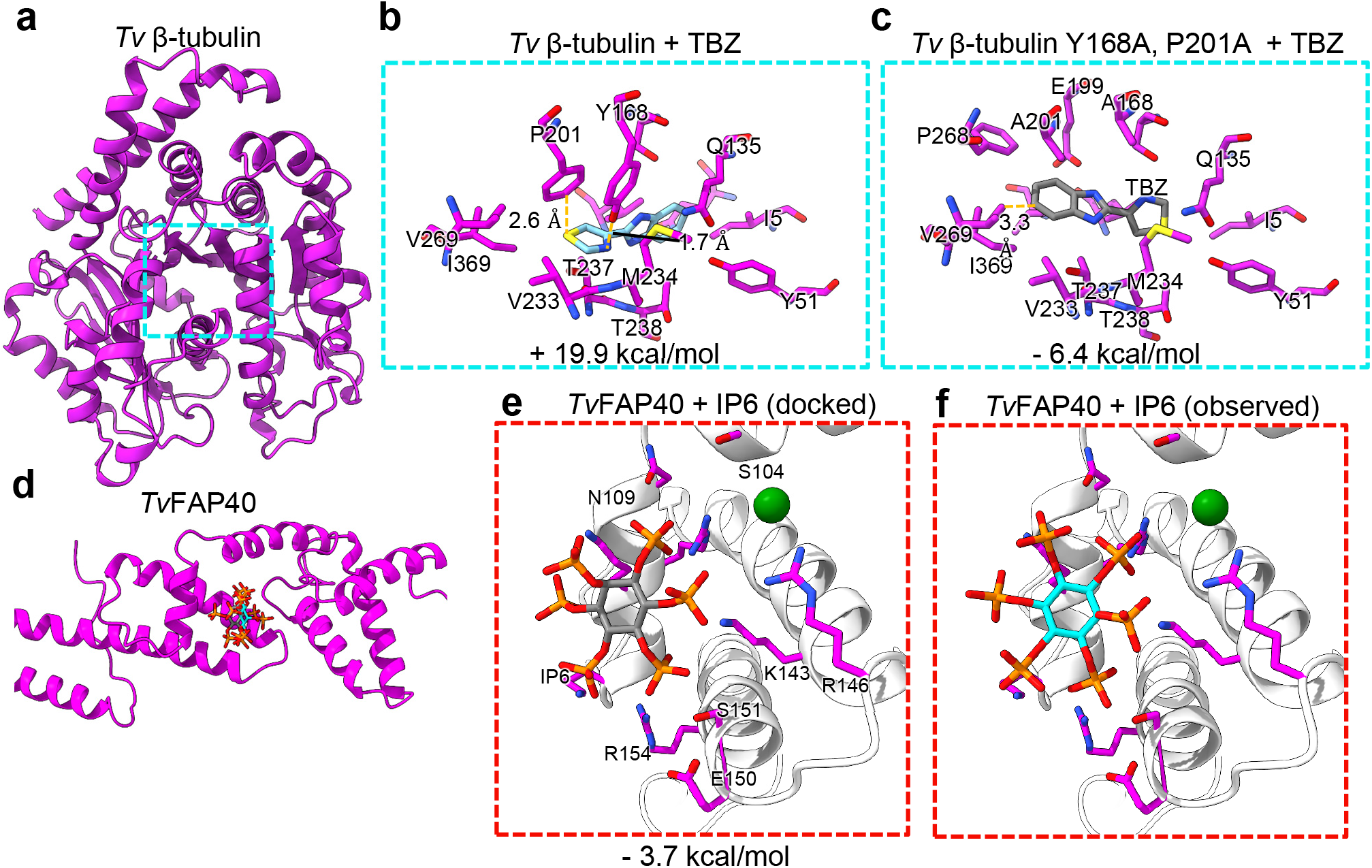
Docking experiments of β-tubulin and *Tv*FAP40. (**a**) Atomic model of β-tubulin with putative BZ drug binding site boxed. **(b)** WT *Tv* β-tubulin with docked thiabendazole (TBZ), fit into putative binding site. (**c**) *Tv* β-tubulin Y168A, P201A mutant with docked TBZ in putative binding site. (**d**) Atomic model of *Tv*FAP40 with putative IP6 binding site boxed. (**e**) *Tv*FAP40 binding pocket with docked IP6. (**f**) *Tv*FAP40 binding pocket with observed IP6.

**Figure S3.**
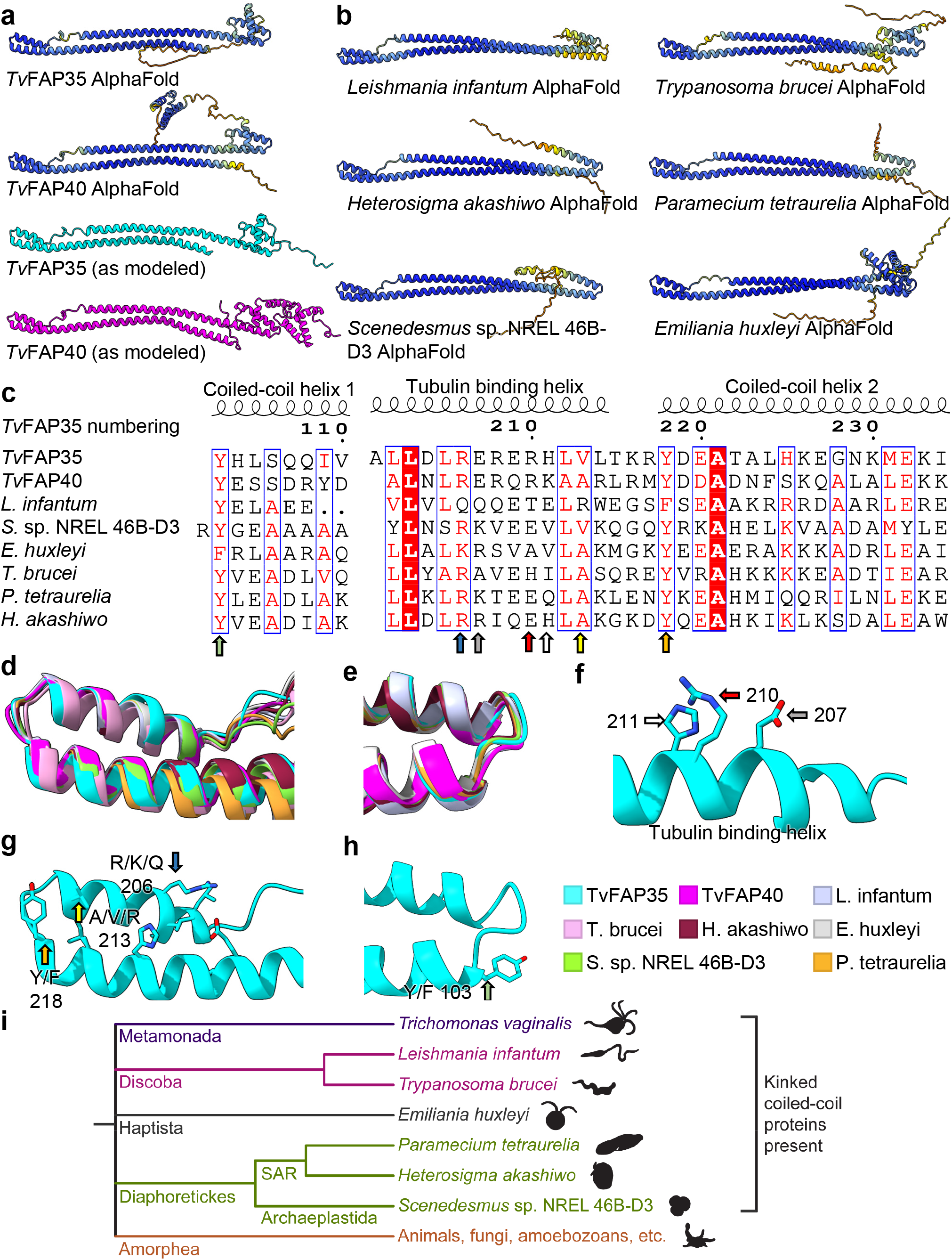
Analysis of *Tv*FAP40 and *Tv*FAP35 and structural homologs. (**a**) AlphaFold-predicted models of *Tv*FAP35 and *Tv*FAP40 (top) colored by AlphaFold confidence interval (blue more confident, red less confident) and their atomic models (bottom) colored in cyan and magenta respectively. **(b)** AlphaFold-predicted structures for structural homologs from selected species, colored by AlphaFold confidence interval. (**c**) Sequence alignment of dimerization and MT binding domain regions from proteins in **a** and **b** aligned to *Tv*FAP35, with conserved residues highlighted and those at the active site indicated with arrows. (**d and e**) ⍺-carbon backbone aligned models from the MT-binding and dimerization domains of the kinked-coiled-coil domains. (**f**) Conserved proteins from **c** shown at their locations at the MT-binding interface on *Tv*FAP35. (**g-h**) Same as **f** but based on both faces of the dimerization domain. (**i**) Phylogeny tree including organisms in which FoldSeek identified similar protein structures.

**Movie S1.**
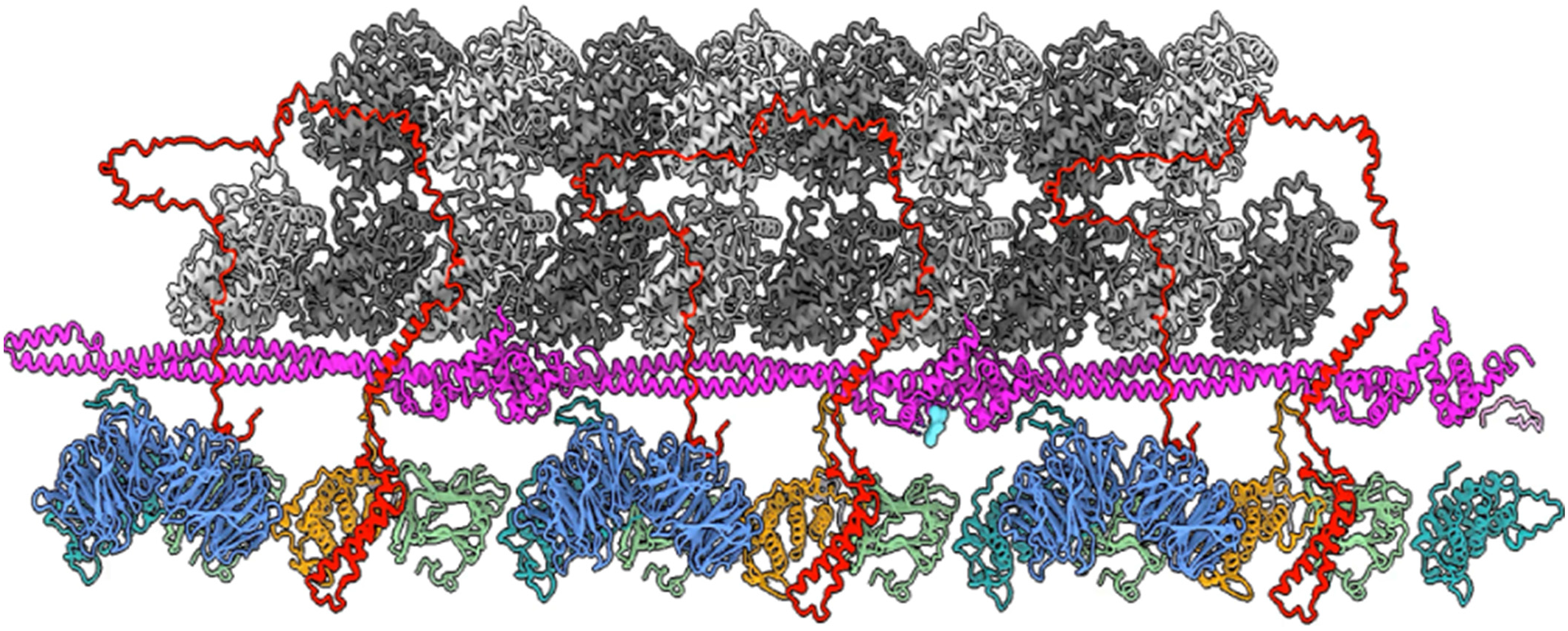
*Tv*FAP40 substrate binding pocket. View flying into putative substrate binding site of *Tv*FAP40. Rotations around the ligand binding site with and without the cryo-EM density.

**Supplementary Table 1.** Cryo-EM data collection.

|  | WT *Tv*-DMT<br>16 nm repeat<br>(EMD-XXXXX)<br>(PDB XXXX) | WT *Tv*-DMT 48<br>nm repeat<br>(EMD- XXXXX)<br>(PDB XXXX) | WT *Tv*-DMT 96<br>nm repeat<br>(EMD- XXXXX)<br>(PDB XXXX) |
| --- | --- | --- | --- |
| **Data collection and processing** |  |  |  |
| Magnification | 81,000 | 81,000 | 81,000 |
| Voltage (kV) | 300 | 300 | 300 |
| Electron exposure (e<sup>-</sup>/Å<sup>2</sup>) | 45 | 45 | 45 |
| Defocus range (μm) | -1.5 to -2.5 | -1.5 to -2.5 | -1.5 to -2.5 |
| Pixel size (Å) | 1.1 | 1.1 | 1.1 |
| Symmetry imposed | C1 | C1 | C1 |
| particle images (no.) | 425,317 | 148,707 | 76,082 |
| Map resolution (Å) | 3.8 | 4.2 | 4.4 |
| FSC threshold | 0.143 | 0.143 | 0.143 |
| Repeat unit (nm) | 16 | 48 | 96 |
| Symmetry imposed | C1 | C1 | C1 |

**Supplementary Table 2.**
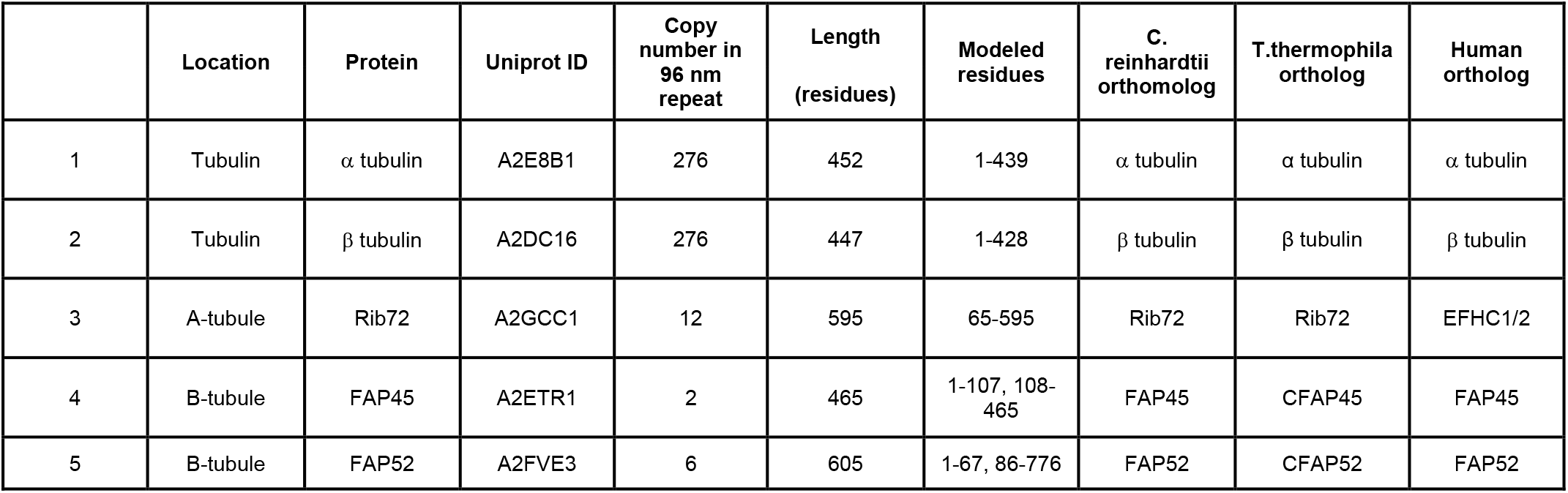
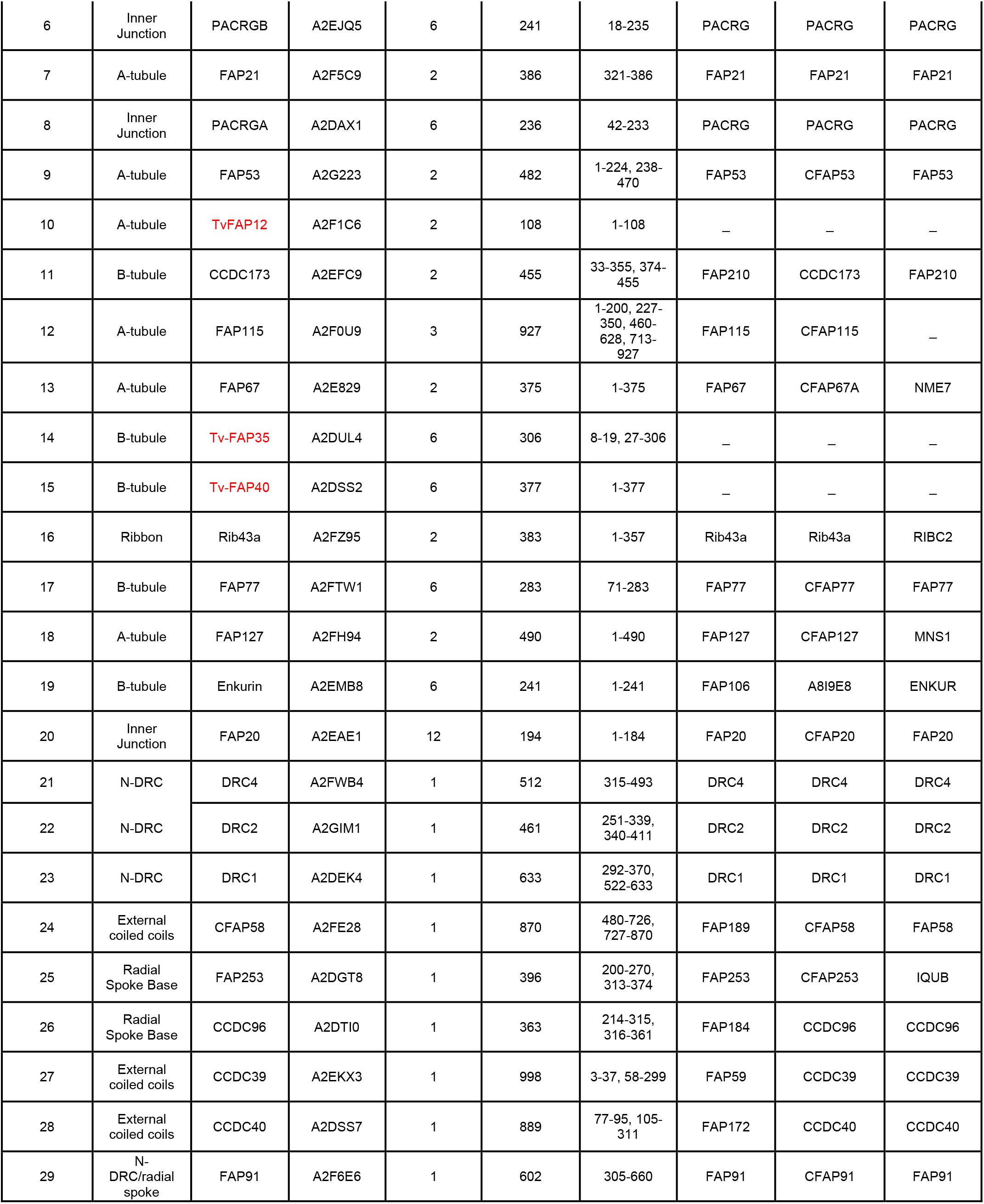
MIPS and MOPS.

